# baltic: the Backronymed Adaptable Lightweight Tree vIsualisation Code

**DOI:** 10.64898/2026.09.20.752947

**Authors:** Barney I. Potter, Karthik Gangavarapu, Sidney M. Bell, María Fernanda Torres Jimenez, Gytis Dudas

## Abstract

For ten years, baltic (Backronymed Adaptable Lightweight Tree vIsualization Code) has been used to make annotated phylogeny figures in molecular epidemiology, including during the West African Ebola epidemic, the Zika epidemic in the Americas, and the SARS-CoV-2 pandemic, as well as other fields. At its core, baltic is a Python library used for the efficient parsing, traversal, manipulation, and visualisation of phylogenetic trees. It is a small library with few dependencies that reads tree formats common in phylodynamic analyses and gives the user leverage to interact with a lightweight tree data structure to produce publication-ready figures with matplotlib. We present baltic v1.0, its first formally released and documented version. baltic reads and writes BEAST Nexus, Newick, and Nextstrain/Auspice JSON, and can process large BEAST posterior tree files in parallel to extract user-defined posterior statistics. The same objects are used for tree manipulation and for plotting in a single script. New to this release: a set of rooting methods (midpoint rooting, rerooting on any branch, and root-to-tip regression); support for reticulate evolution, with reassortment and recombination edges; and composite figures that combine a tree with other data, such as Müller plots, skygrid plots, tanglegrams, and plots connecting trees to maps. The release includes a documentation site with an API reference, tutorials, and a matplotlib-style gallery of worked examples.

## Introduction

### Modern phylogenies are (too) big

The scale of modern sequence data has transformed phylogenetics from a field limited by data availability into one increasingly constrained by interpretation. For example, since the beginning of the COVID-19 pandemic, over 15 million SARS-CoV-2 genomes have been sequenced and shared on sequence databases. Phylogenetic trees remain the central abstraction for reasoning about evolutionary relationships and population processes, but they are also an inherently difficult visual medium. At their core, modern phylogenetic visualisations are at best pseudo-two-dimensional: typically one of two visualisation dimensions is sacrificed to ensure taxa do not overlap and to convey the nested hierarchy of relationships between taxa, while the other visualisation dimension is commonly used to depict the amount of genetic or temporal distance on branches of the tree. In combination with increasingly more taxa and metadata, trees rapidly become dense, label-heavy, and uninterpretable at first glance. This is further exacerbated by sophisticated analyses that include phylogenetic uncertainty, ancestral state reconstruction, novel data structures (*e*.*g*. multitype trees, reassortment networks), as well as new formats (*e*.*g*. `Nextstrain`’s `Auspice` JSON). We aim to address these shortcomings via novel visualisation techniques which we implemented in an updated Python software package. Here we present `baltic`, the **b**acronymed **a**daptable **l**ightweight **t**ree v**i**sualisation **c**ode, a Python (*≥*3.5) module designed for parsing and manipulating phylogenetic trees which leverages `matplotlib` (*≥* 2.0.0) [1] as the visualisation engine. `baltic` focuses on user friendliness, thoughtful static data visualisation and modern phylogenetic analysis outputs. While `baltic` has seen some use in the past [2–181], here we present the latest version 1.0 with an improved API and newly implemented features.

### baltic philosophy and gallery

When writing `baltic`, we were heavily inspired by `matplotlib`, which `baltic` uses for rendering. Like `matplotlib`, `baltic` providers users leverage and flexibility to construct complex visualisations out of simpler elements. For example, one of the most argument-rich functions in `matplotlib`’s code base is used for drawing box-and-whiskers plots, arising out of the need to give users aesthetic control over each sub-element of the visualisation. While `baltic` also implements some field-specific advanced visualisations as single functions (*e*.*g*. exploded trees or state-collapsed trees), our goal is to strike a balance between utility and functional simplicity. To this end, we follow `matplotlib`’s example and rather than trying to foresee every possible `baltic` use case, we built a gallery of examples using `baltic` to showcase how various `baltic` functions can be used together to produce more advanced visualisations (*e*.*g*. Fig. 1). This gives users a solid starting point and complete artistic and study-specific control over the users’ visualisations.

**Fig. 1.**
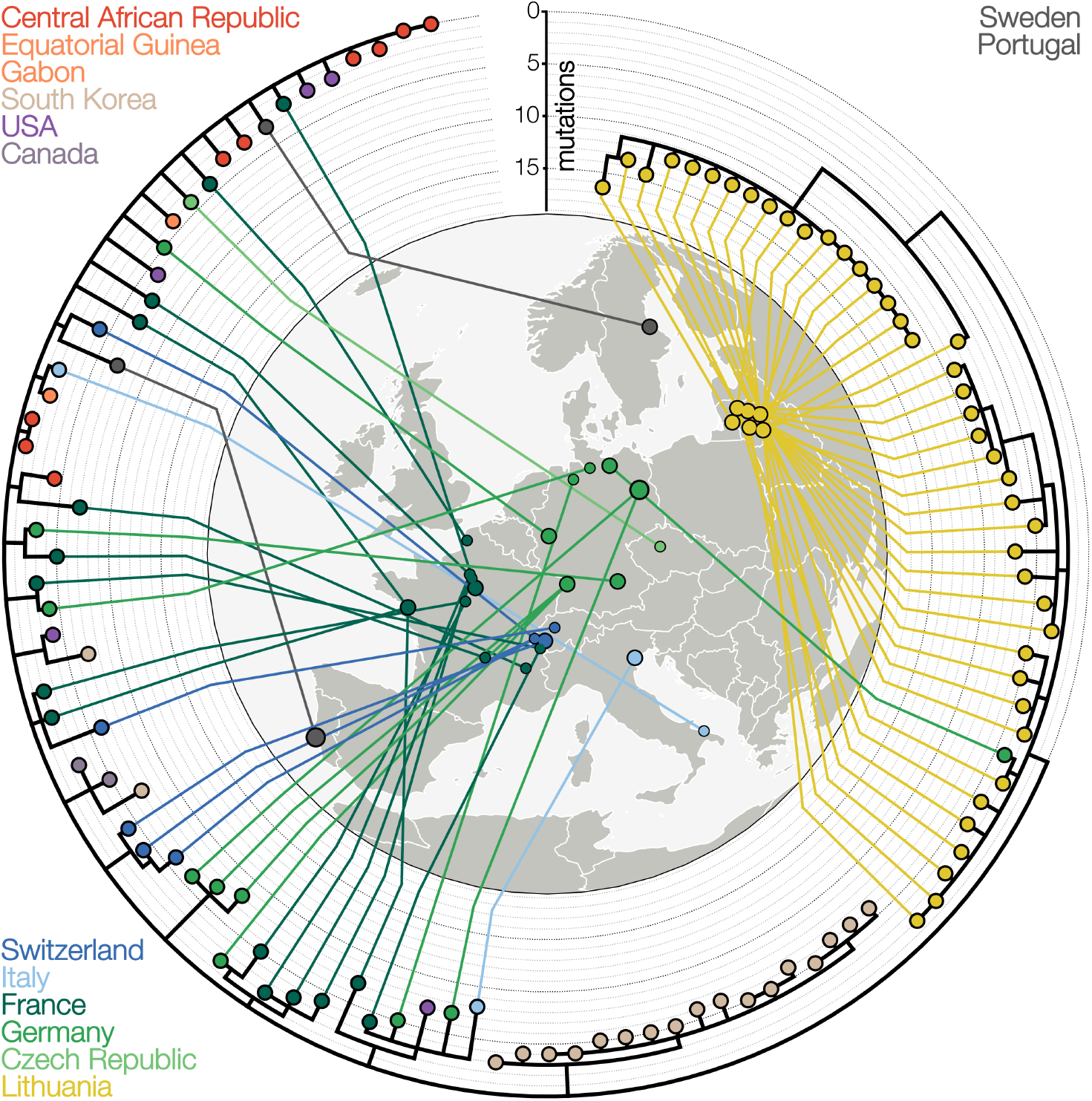
An inward-facing circular tree around a map of Europe adapted from [6] showing a maximum likelihood tree of SARS-CoV-2 lineage B.1.620, first recognised in Lithuania, and connected to its sampling locations and coloured according to country.

## Implementation(s)

### baltic core

`baltic` parses phylogenetic trees supplied in Newick, Nexus, or `Nextstrain`/`Auspice` JSON format [182] into a common in-memory representation. For Newick- and Nexus-derived strings, `make tree` performs a single left-to-right pass over the tree string with a cursor index, incrementally matching tokens for parentheses (new internal nodes), tip labels (both plain and `BEAST`-style integer-encoded taxa), branch lengths, and annotation comments (*e*.*g*. `[&posterior=1.0,…]`), which are parsed into a per-branch `traits` dictionary; reticulate edges denoting recombination, reassortment, or conversion events are recognized via `#`-prefixed labels and linked to their destination once both ends have been seen. The `baltic.io` module provides format-specific wrapper functions: `load newick` and `load nexus` locate the tree string within a file (the latter also resolving Nexus `Translate` blocks that map numeric tip codes to full taxon names) and can extract tip sampling dates from taxon labels via a user-supplied regular expression, while `load JSON` walks an `Auspice` v2 JSON tree recursively (`make tree JSON`), remapping `Nextstrain`’s `node attrs`/`branch attrs` fields onto the same internal attributes used by `baltic` Newick/Nexus parsers so that all three input formats converge on one tree object.

Every parsed tree has a `treeType` of either `“divergence”` or `“time”`, which governs how branch lengths are interpreted: in a divergence tree, lengths represent substitutions (or another divergence unit) and a pre-order traversal (`traverse tree`) accumulates them into a `height` measured from the root, whereas in a time tree the same accumulated height is additionally mapped onto calendar time (`absoluteTime`) using either a most recent sampling date (`set absolute time`) or dates parsed directly from tip labels. Structurally, `baltic` stores each branch as an instance of a class hierarchy rooted in the `BranchLike` superclass, which holds attributes common to every branch (incoming `length`, `height`, `absoluteTime`, `parent`, a `traits` dictionary of parsed annotations, and plotting coordinates). `BranchLike` is subclassed into `Leaf` (terminal taxa), `Node` (internal ancestors, which additionally track their `children` and the set of descendant tip names), `Reticulation` (non-vertical edges representing reticulate events, linked to a `target` branch elsewhere in the tree), and `Clade` (a placeholder representing a collapsed monophyletic subtree for compact display).

These branch objects are held collectively by a `Tree` object, which stores a flat list of all `BranchLike` objects (`Objects`), a reference to the `root`, the cursor (`curNode`) used during parsing, and exposes traversal, sorting, statistics, and plotting methods that operate uniformly across the branch-object hierarchy regardless of input format.

### Python functions in baltic

Much of `baltic`’s flexibility comes from its canonical use of, and default design for Python functions as arguments (particularly Python’s `lambda` functions). Most of `baltic`’s API interfaces, be they for selecting taxa for plotting or describing a color mapping, expect functions (typically taking just the argument `k`, a `baltic.branchLike` object) as their argument. The generous use of this Python feature allows users great flexibility in the specific implementations they choose, allowing both very simple functions that only return a single value on any input (*e*.*g*. a colour function that only returns “black”), or the expansion of more complex mappings into complex functions.

### baltic format interoperability

`baltic` supports several file formats commonly used in phylogenetics, and though newick and nexus formats have been standard for many decades, their full range of flexibility has not always been supported by downstream software. For example, the comment format of nexus files is widely employed within the `BEAST` [183] ecosystem to encode information like branch-specific molecular clock rates and ancestral state reconstruction but this is not always supported by all phylogenetic tree parsers. Extensions beyond strictly clonal trees like ancestral recombination graphs, reassortment networks and other reticulate data structures are accommodated by even fewer parsers. Though newick and nexus file formats remain the *de facto* community standard, there are competing formats. For example, the `Auspice` JSON format developed and used in the `Nextstrain` ecosystem is often employed in genomic epidemiology (*i*.*e*. public health) contexts.

`baltic` currently supports newick, nexus, and `Auspice` JSON file formats as inputs, as well as the extended `newick` format as seen in the `BEAST2` [184] ecosystem and implemented in `IcyTree` [185]. Of these, the ability to handle the `Auspice` JSON format is of particular note, as few phylogenetic parsers are currently able to handle it. To the best of our knowledge these are `treeio` [186] for R, and `Nextstrain`’s `Auspice` with `taxonium` [187] for javascript-based web visualisation but nothing, to our knowledge, for Python. Likewise, parsing reticulate data structures remains rare, with `PhyloNetworks.jl` [188] (Julia) and `IcyTree` (javascript/web) being the few examples, and nothing available, to our knowledge, in Python currently.

### Standard visualisations

Users of `baltic` have access to all the typical phylogenetic visualisation approaches. For large subtrees, `baltic` offers the possibility of collapsing clades into narrower triangles to conserve space. For cases where some internal branches are not needed, *e*.*g*. polytomies resolved into arbitrary bifurcations (either with zero or miniscule branch length) or low statistical support, `baltic` allows users to collapse these according to custom filtering functions. If a user is more interested in specific clades, `baltic` offers the ability to extract subtrees descended from any node. `baltic` also allows users to reroot (substitution space, *i*.*e*. `treeType=‘divergence’`) phylogenetic trees, and implements root-to-tip regressions for measurably evolving populations. Examples of subtree extraction, collapsing clades, rerooting, and padding nodes (see “Padded nodes and tanglegrams” section) are shown in Fig. 2.

**Fig. 2.**
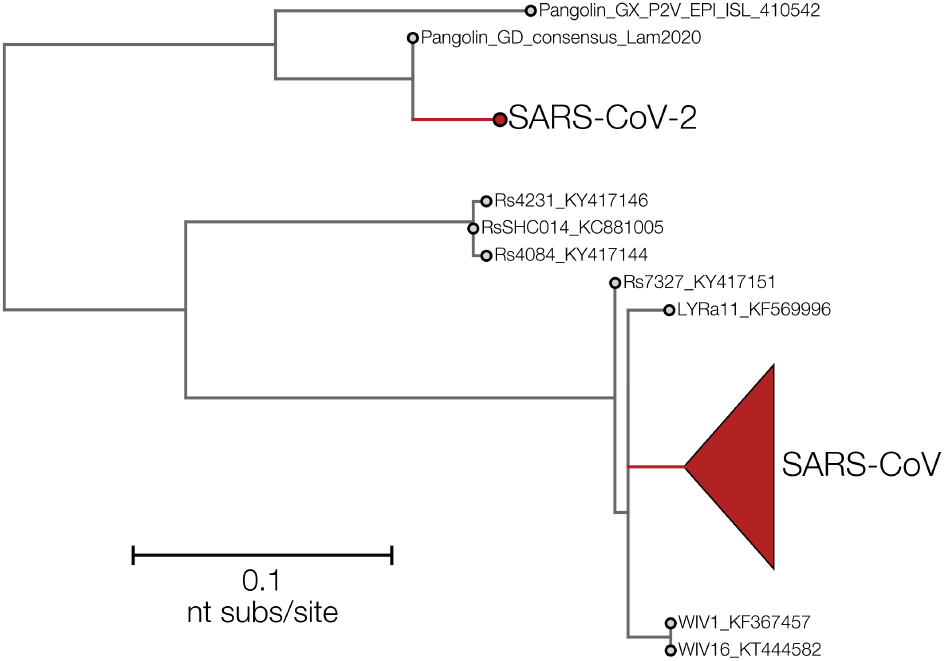
A maximum likelihood phylogenetic tree of betacoronaviruses from [189]. SARS-CoV (collapsed clade) and SARS-CoV-2 are highlighted in red, with extra vertical space added around both using baltic’s padNodes feature.

### Exploded trees

During the 2013–2015 West African Ebola virus epidemic, Philippe Lemey (KU Leuven) developed a visualisation for Ebola virus trees annotated with reconstructed geographic regions [10]. The visualisation depicted the time span between the arrival of an Ebola virus lineage to a new region and the sampling of that introduction’s last descendant in the same region, ignoring lineages that migrated elsewhere. This approach conveys information about the number, size (in terms of sequenced genomes) and duration of each sub-outbreak. This idea was implemented in `baltic` using conditional tree traversals - each branch whose parent is in a different discrete state initiates a tree traversal conditioned on each child branch remaining in the new location, with subtrees resulting in tip-less subtrees being discarded. The resulting state-specific subtrees are grouped together and displayed, often with the pre-introduction state visually indicated with colour (Fig. 3A). This rearrangement of the full tree into blocks based on discrete traits allows the impact of each trait state’s effect on the fate of introduced lineages to be immediately interpretable. Andrew Rambaut (University of Edinburgh) coined the term “exploded trees” for this visualisation and it has been used to define “transmission lineages” [190] during the COVID-19 pandemic and later implemented in the `Nextstrain` ecosystem under the “Explode tree by” option.

**Fig. 3.**
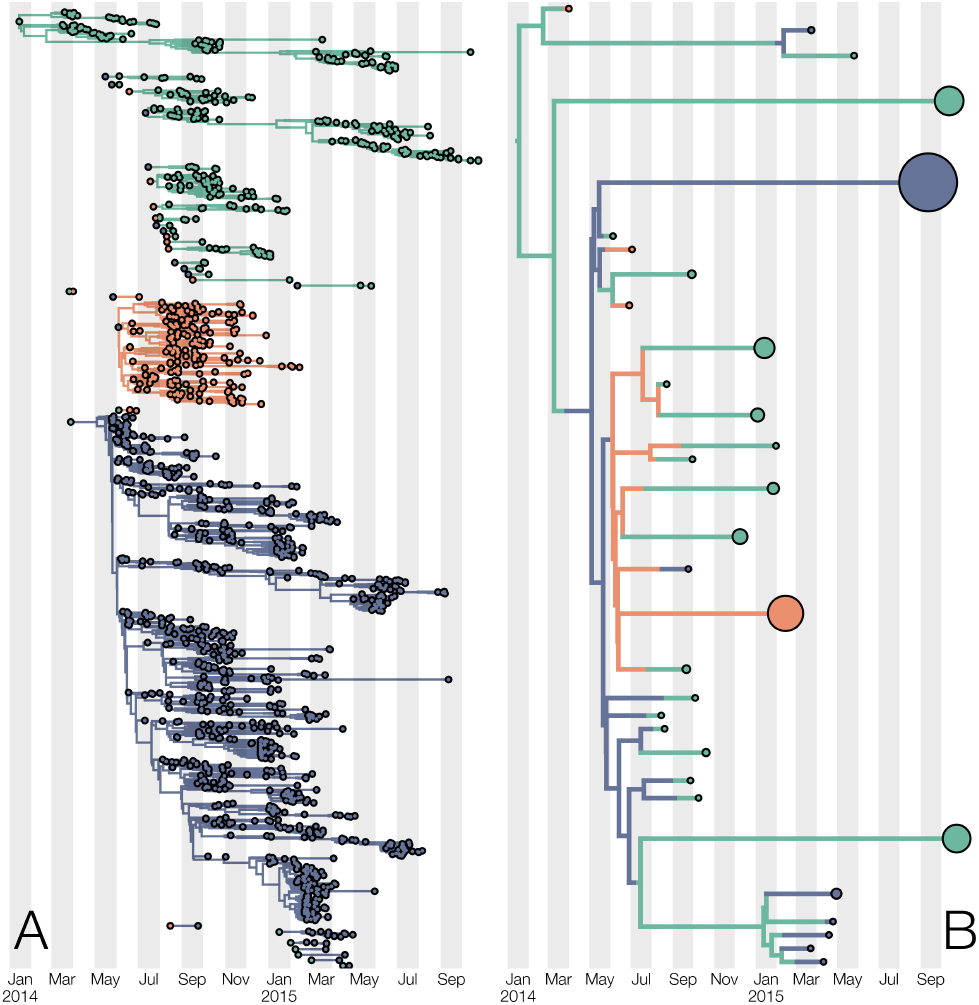
**A**) An “exploded tree” visualisation derived from a molecular clock tree of 1610 Ebola virus genomes from the 2013-2015 West African Ebola virus epidemic from [10]. Each subtree begins with an international migration between the three most affected countries during the epidemic (Guinea in green, Sierra Leone in blue, Liberia in red) and tracks lineages for as long as they stay in the country of introduction. **B**) A “state-collapsed” visualisation derived from the same original tree as A). Rather than displaying entire subtrees (A), it preserves the phylogenetic position of the earliest tip descended from an international introduction, whose “tip” date is adjusted to indicate when the last sampled descendant of the introduction was sampled and circle size is scaled according to how many tips in total descend from the introduction. As before, Guinea is shown in green, Sierra Leone in blue, and Liberia in red.

### State-collapsed trees

While the exploded tree visualisation merely rearranges an existing tree topology by sacrificing branches that have state changes and retaining relationships within the same state, there is a more compact approach. Initially developed around 2017 for the visualisation of the 2013–2015 West African Ebola virus epidemic and never used in a publication, the method involves partitioning the tree into stretches of continuous evolution under a given trait state following a migration. This is done by traversing the tree from the root with a heritable unique label which switches to a new label if a child branch switches to a new trait state. As a result, these assigned labels partition the tree into groups of related branches that represent continuous evolution in the new trait state. Then, for each unique label the earliest sampled tip is chosen to represent the entirety of that label, its height is adjusted to correspond to the last sampled tip with the same label, the number of tips with that label is represented by the circle size of the representative tip, and all non-representative tips are removed from the tree while preserving their phylogenetic relationships (see next sections). We call this visualisation “state-collapsed trees” (Fig. 3B), as it yields a compact visualisation of the history of trait transitions with the final number of tips equal to the number of transition events. Due to the pruning procedure state-collapsed trees yield multitype trees (with node objects that have a single descendant branch) that retain information about the timing (for time trees) of transition events as well. State-collapsed trees are conceptually close to phylotype maps from `PhyloType` [191], and inter-cluster transition trees from `EvoLaps2` [192] but yield phylogenetic data structures (multitype trees) that can still be imported by common phylogenetic software like `FigTree` [193], though trees with reconstructed ancestral states at internal nodes are arguably the most impactful use case.

### Reduced trees, rerooting, and processing posterior distributions

Like many other phylogenetic parsing, analysis, and visualisation packages, `baltic` allows users to reroot non-molecular clock trees (translated from `BioPython`’s Phylo module), drop individual tips, and depict trees in circular and unrooted configurations. Owing to its genomic epidemiology roots, `baltic` also allows users to automatically infer or manually assign tip dates in absolute time as well as their uncertainty (*e*.*g*. when only the sample year but not full date of collection is known), and infer the root of the tree that optimises (via sum of squares, *r*^2^ or correlation coefficient) a root-to-tip regression. Similarly, `baltic` can process posterior sets of trees in `BEAST` nexus format to extract various posterior statistics of interest, *e*.*g*. common ancestor dates (TMRCAs) or discrete trait probabilities along a single path between a tip and the root. Posterior tree set handling is done via a submodule of `baltic` called `samogitia` where users can use custom functions for processing each posterior realisation of a tree and return processed outputs to a standard Bayesian phylogenetic trace/log file.

### Padded nodes and tanglegrams

Given the ever-growing size of phylogenetic trees, `baltic` natively implements explicit padding around specified branches. This is similar to `ggtree`’s [194] (R) `scaleClade` function, though in the `baltic` implementation users provide a dictionary (`padNodes`) where keys are branch objects and values are `int`s or `float`s that specify the amount of vertical space to be added around the specified branch (in `baltic`, each tip receives 1 unit of vertical space around it by default). `baltic` also supports tangled chain visualisations, where two or more rooted trees are displayed in series with tips representing the same sample in different trees are connected with lines to highlight phylogenetic incongruence (Fig. 4A). However, `baltic` provides the flexibility to easily extend this visualisation type to unrooted trees by providing users access to tree plotting coordinates (Fig. 4B). We also provide the option to “untangle” (reduce connecting line crossing) trees prior to plotting tangled chains, though we believe it to be misleading since trees can have tips positioned at the same y-axis coordinates (and thus have non-intersecting tanglegram lines) whilst not being phylogenetically congruent.

**Fig. 4.**
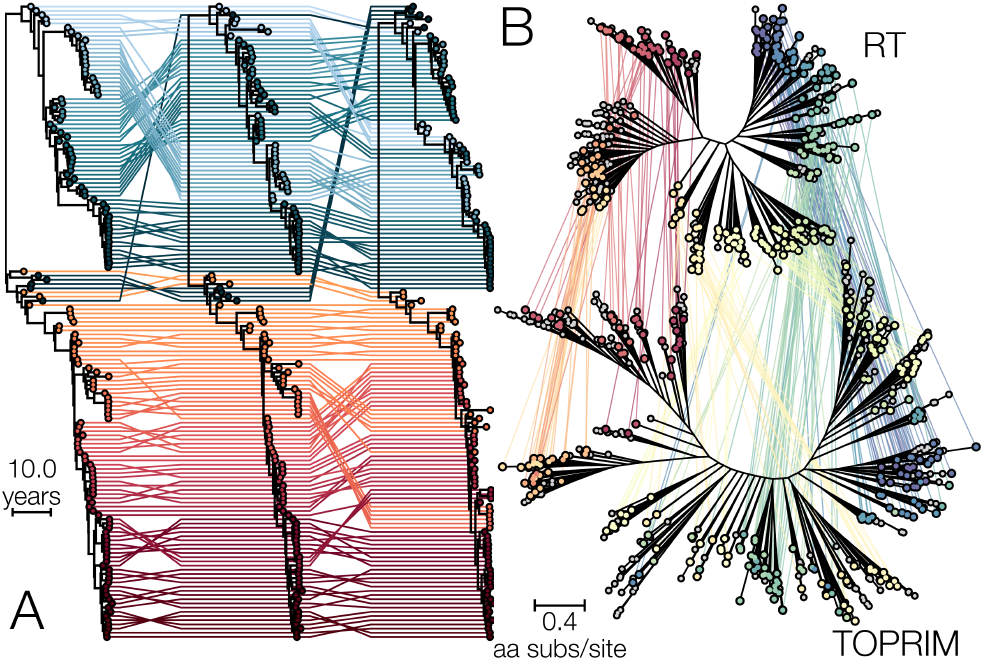
**A**) A tangled chain of influenza B virus PB1, PB2 and HA segment time trees (in that order from left to right) from [18], down-sampled three-fold and with coloured lines connecting sequences derived from the same genome across all trees. Blue lines indicate Yamagata line-age sequences (in HA tree), red lines are Victoria lineage (in HA tree) with different shades indicating the y-axis position of each tip in the PB1 tree. **B**) An unrooted tanglegram of reverse transcriptase (RT) and topoisomerase-primase (TOPRIM) domains from retron types I-C, XI, and XII connecting domains derived from the same protein from [150]. Tip circles and connecting lines are coloured according traversal order from blue to red.

### Müller plots

Müller plots are a unique and uncommon visualisation technique that preserves phylogenetic information and depicts absolute or relative genotype frequencies through time [196]. It was first used to intuitively explain the concepts of Müller’s ratchet, clonal interference and the advantages of recombination in generating fit genotypes. Typically, the x-axis of Müller plots conveys the passage of time, the y-axis shows genotype frequencies and the hierarchical relationships between genotypes is depicted as descendant genotype frequencies emerging from the middle of the parental genotype frequency. There are relatively few software packages for doing Müller plots, with `ggmuller`, `MullerPlot`, and `EvoFreq` [197] being notable examples in the R ecosystem, and `pyfish` [198] and `muller` in the Python ecosystem. The `Augur` (Python) package, the back-end of `Nextstrain` analyses, is a unique example, where clade frequencies for individual phylogenetic trees are computed via nested logistic smoothing of temporal observations (tips in the tree). While `Augur` infers the frequency trajectories that are the basic building block of Müller plots, `Auspice`, the front-end part of `Nextstrain` doesn’t display them as Müller plots in the strictest sense. In `Auspice`’s version of Müller plots simpler stacked area plots. In `baltic` we offer users an option for plotting frequencies such that the starting y-axis coordinates of descendant genotypes begin in the middle of the parental genotype frequency, thus preserving information about phylogenetic relationships. This stylistic choice is made via the boolean `Muller` argument of `baltic`’s `plot Muller` function (`True` for strict Müller diagrams with descendant frequencies emerging from the ancestor frequency and `False` for `Auspice`-style plots) in `baltic`.

In `baltic`, we have reused `Augur`’s code to compute clade frequencies but modified it so observations (dates of tips) can be assigned directly to nodes in the tree. By doing this, we enable the user to compute and depict Müller plots with millions of observations while maintaining a minimal underlying phylogenetic data structure. The typical use case we envision is taking an abstract lineage tree (*e*.*g*. pango nomenclature for SARS-CoV-2 [199]) and annotating its branches with sample collection dates of each lineage. An example of this is shown in Fig. 5, which depicts the relationships and frequencies of over 1 million SARS-CoV-2 genomes from UK’s SARS-CoV-2 genomic surveillance programme.

**Fig. 5.**
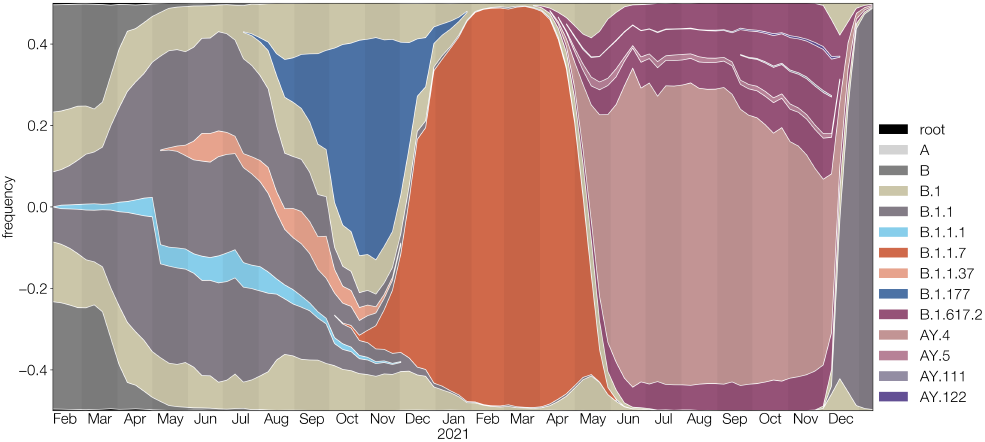
A Müller plot of SARS-CoV-2 derived from more than 1 million genomes using data from [195]. The Müller plot depicts both the relative frequencies of multiple pre-specified SARS-CoV-2 lineages (indicated by colour, legend to the right) and abstractly depicts their phylogenetic relationships derived from pangolin lineage labels.

## Conclusion

`baltic` is a lightweight and flexible Python tool for parsing, analysing, manipulating and visualising phylogenetic trees. By exposing much of the underlying (*e*.*g*. traits) and derived (*e*.*g*. plotting coordinates) attributes of the phylogenetic data structure, we give users the freedom to analyse, manipulate and process phylogenetic trees, and build their own publication-ready static figures. To help new users we provide a useful starting point with extensive tutorials and gallery. We encourage first time users to provide feedback that could improve `baltic`’s usability and advanced users to request support for non-standard phylogenetic data structures or visualisations.

## Acknowledgments

Over the years, many people contributed to `baltic` directly or indirectly by regular use and inadvertent testing. In particular, we’d like to thank Anderson Brito, Verity Hill, Gage Moreno, Louise Moncla, Edyth Parker, Miguel Paredes, Jesse Bloom, John Huddleston, Nicola Müller, Áine O’Toole, Jonathan Pekar, Allison Black, Ifeanyi Omah, Matthew Scotch, Raphäelle Klitting, Alvin X. Han, Praneeth Gangavarapu, and many others. A special thanks goes to Andrew Rambaut and Trevor Bedford for support, advice, ideas, and dedication to the craft of data visualisation in phylogenetics.

GD acknowledges funding from the Research Council of Lithuania, Lithuania under the EMBO Installation Grant programme under grant No 5305.

## Materials and Methods

### Data availability

Figure 1 is a simplified tree from [6]. GISAID accessions from a study of SARS-CoV-2 lineage B.1.620 [6] were recovered from a publicly available tree file https://github.com/evogytis/B.1.620-in-Europe/blob/main/data/trees/latest_continent_mcc.tre). All accessions were downloaded from GISAID (for acknowledgments see EPI SET https://epicov.org/epi3/epi_set/260918gh), aligned using `NextClade` CLI [200], hand-picked for quality, and the maximum likelihood phylogenetic tree recovered using `RAxML` v.8.2.11 [201] under a GTR [202] + Γ_4_ [203] model.

Figure 2 is derived from a study into ACE2 binding of coronaviruses [189]. The tree used was publicly available at https://github.com/jbloomlab/SARSr-CoV_homolog_survey/blob/52db008d423a17d622950d6311fd89392cdc5e1d/RBD_ASR/RBD_rm_RaTG13/ASR/tree.newick.txt. Prior to visualisation, the clade descending from the common ancestor of SARS-CoV and SARS-CoV-2 was extracted using the `subtree` method of the `baltic.Tree` class, midpoint rooted and the same `subtree` method called again on the common ancestor of SARS-CoV and SARS-CoV-2.

Figure 3A is a reproduction of Figure 4 from [10], and both it and Figure 3B are derived from a publicly available tree at https://github.com/ebov/space-time/blob/master/Data/Makona_1610_cds_ig.GLM.MCC.tree.

Figure 4A depicts three of eight segment trees of influenza B virus from a study into reassortment dynamics between Victoria and Yamagata lineages of this virus [18]. The trees are available at https://github.com/evogytis/fluB/blob/master/data/mcc%20trees/InfB_PB1t_ALLs1.mcc.tre, https://github.com/evogytis/fluB/blob/master/data/mcc%20trees/InfB_PB2t_ALLs1.mcc.tre, and https://github.com/evogytis/fluB/blob/master/data/mcc%20trees/InfB_HAt_ALLs1.mcc.tre.

Figure 4B is a reproduction of Supplementary Figure 1 from [150], showing the phylogenetic trees of reverse transcriptase (RT) and topoisomerase-primase (TOPRIM) domains of retron Eco2 and its relatives. Original tree files and annotations are available from https://data.mendeley.com/datasets/v9b289v6ct/1.

Figure 5 is derived from data on SARS-CoV-2 lineage dynamics in the UK [195] and is based on raw data available at https://github.com/qinqin-yu/sars-cov-2_genetic_drift/blob/main/data/metadata/cog_england_2022-01-16_metadata.csv.

All data and code to reproduce figures provided here are available at https://github.com/phylo-baltic/baltic-v1-manuscript. The code for `baltic` is available at https://github.com/evogytis/baltic.

### User resources

We provide documentation for `baltic` at https://baltic.readthedocs.io/en/latest/index.html. Recognising that `baltic`’s strengths lie in combining multiple features in non-trivial ways, we also provide a gallery and associated code for advanced visualisations at https://phylo-baltic.github.io/baltic-gallery/.

### Large language model use disclosure

At various points during the development of baltic v1.0 we used Codex (ChatGPT 5.4, 5.5, and 5.6 (High)) and Claude to fix bugs, optimise and implement new functions, and preparation of documentation.

**Table 1:**
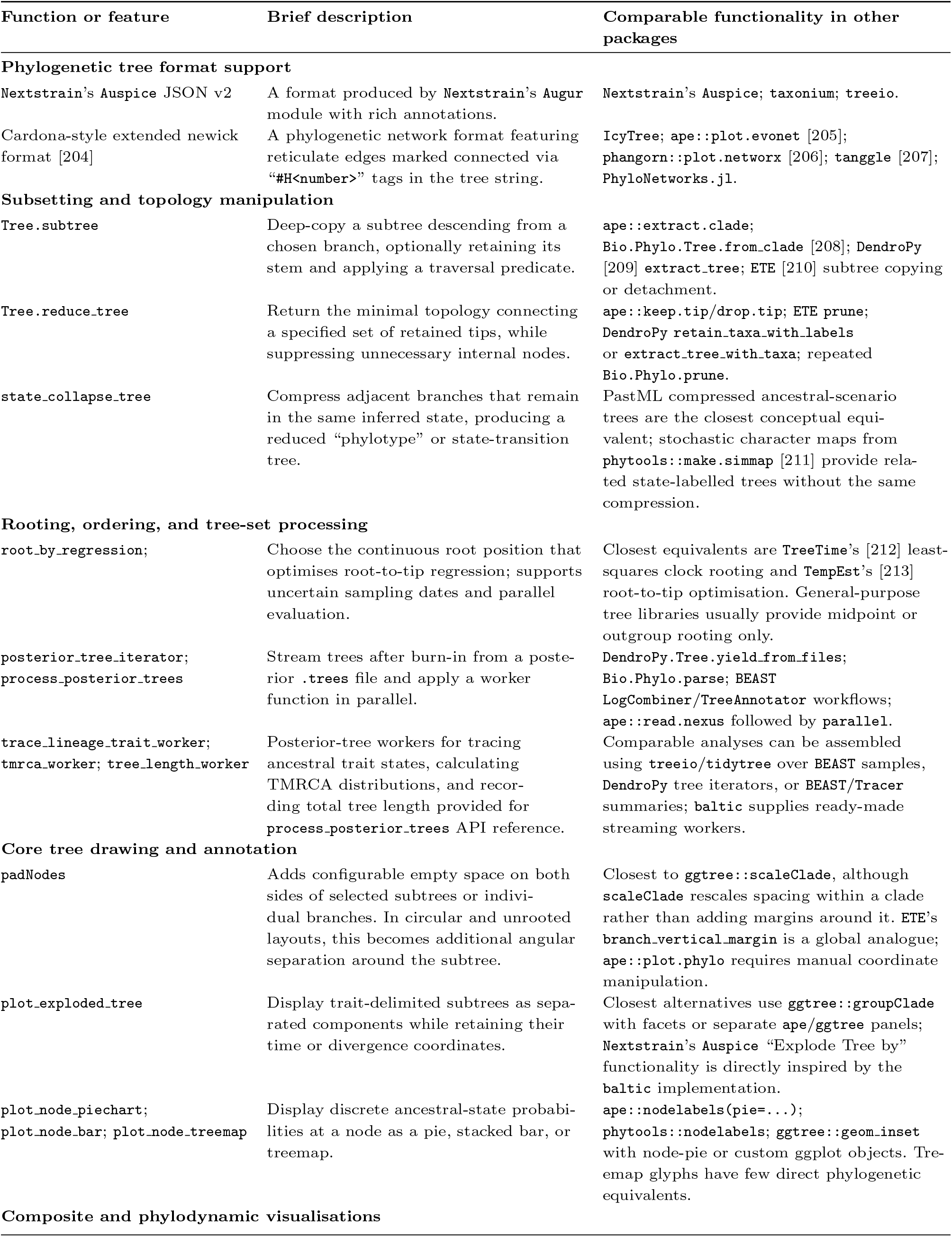
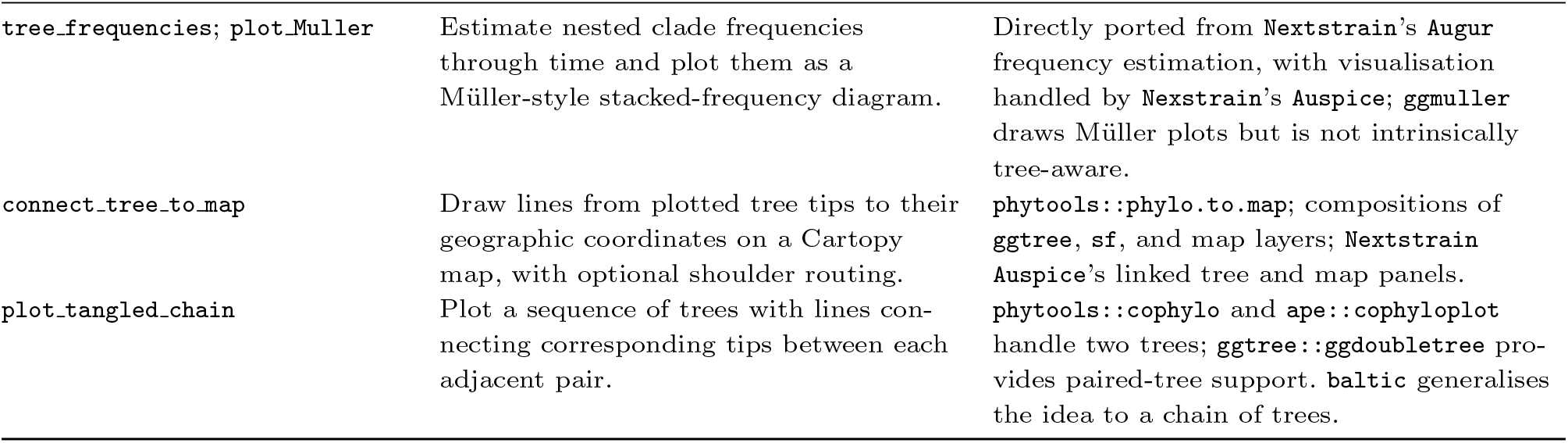
Highlights of baltic’s functions and features with comparable functionality in other phylogenetic software.

